# Antigen non-specific CD8^+^ T cells accelerate cognitive decline in aged mice following respiratory coronavirus infection

**DOI:** 10.1101/2024.01.02.573675

**Authors:** Katie L. Reagin, Rae-Ling Lee, Loren Cocciolone, Kristen E. Funk

## Abstract

Primarily a respiratory infection, numerous patients infected with SARS-CoV-2 present with neurologic symptoms, some continuing long after viral clearance as a persistent symptomatic phase termed “long COVID”. Advanced age increases the risk of severe disease, as well as incidence of long COVID. We hypothesized that perturbations in the aged immune response predispose elderly individuals to severe coronavirus infection and post-infectious sequelae. Using a murine model of respiratory coronavirus, mouse hepatitis virus strain A59 (MHV-A59), we found that aging increased clinical illness and lethality to MHV infection, with aged animals harboring increased virus in the brain during acute infection. This was coupled with an unexpected increase in activated CD8^+^ T cells within the brains of aged animals but reduced antigen specificity of those CD8^+^ T cells. Aged animals demonstrated spatial learning impairment following MHV infection, which correlated with increased neuronal cell death and reduced neuronal regeneration in aged hippocampus. Using primary cell culture, we demonstrated that activated CD8^+^ T cells induce neuronal death, independent of antigen-specificity. Specifically, higher levels of CD8^+^ T cell-derived IFN-γ correlated with neuronal death. These results support the evidence that CD8^+^ T cells in the brain directly contribute to cognitive dysfunction following coronavirus infection in aged individuals.

**eTOC summary:** Using a murine model of respiratory coronavirus infection, we show that aging amplifies post-infectious cognitive dysfunction due to activated CD8^+^ T cells that secrete IFN-γ in the brain. These data provide evidence that CD8^+^ T cells in the brain negatively impact post-infectious cognitive function.

## Introduction

Increasing evidence correlates exposure to viral infections with cognitive dysfunction and neurodegenerative diseases(Lotz et al., 2021). Neurotropic viral infections such as Zika virus (Figueiredo et al., 2019; Oh et al., 2017), West Nile virus (WNV)(Fulton et al., 2020), and herpes simplex virus (De Chiara et al., 2019) have long been associated with post-infectious cognitive sequelae attributed largely to immune cell infiltration and inflammation within the brain. However, recent data suggest that exposure to viral respiratory pathogens, including influenza(Jurgens et al., 2012), respiratory syncytial virus(Andrade et al., 2022), and SARS-CoV-2(Ellul et al., 2020), may cause neurologic symptoms and post-infectious cognitive decline(Damiano et al., 2021). Prior exposure to influenza highly correlates with onset of neurodegenerative disorders including Alzheimer’s disease, Parkinson’s disease, and multiple sclerosis(Levine et al., 2023); however, whether other respiratory viral infections, including SARS-CoV-2, may contribute to cognitive decline remains unclear. While SARS-CoV-2 manifests primarily in the lower respiratory tract as viral pneumonia, 36% of patients develop neurologic symptoms of disease including fatigue, myalgia, headache, anxiety and depression(Vanderheiden and Klein, 2022), some of which persist long after resolution of acute viral infection(Taquet et al., 2021), known as “long COVID.” Long COVID was initially described in patients with severe infections, but recent reports have described these symptoms even in patients with relatively mild illness(Méndez et al., 2022). Although there is some evidence of direct central nervous system (CNS) infection by SARS-CoV-2(Stein et al., 2022), it is not clear whether symptoms of long COVID are due to viral invasion per se or may result from neuroinflammatory responses to the virus(Klein, 2022). SARS-CoV-2 infection is characterized by heightened production of inflammatory cytokines including IL-1β, IL-6, and TNF in the plasma(Arunachalam et al., 2020), as well as CNS infiltration of antiviral immune cells, including CD8^+^ T cells(Schwabenland et al., 2021), which can contribute to post-infectious cognitive disease(Reagin and Funk, 2022). Therefore, antiviral inflammation may drive post-infectious cognitive sequelae in long COVID patients. With more than 600 million people worldwide who have survived COVID acute infection(Statista), it is critical that we understand the biological basis of post-acute sequelae, including neurologic dysfunction.

Advanced age is a significant risk factor for severe outcomes of viral infection, including SARS-CoV-2. Individuals over the age of 65 display increased incidence of severe disease(Booth et al., 2021; Abul and Leeder, 2023), risk for re-infection with SARS-CoV-2(Michlmayr et al., 2022), and incidence of long COVID(Sullivan and Fischer, 2021), as well as neurologic complications(Sullivan and Fischer, 2021). This is likely due to age-associated changes in the antiviral immune response, which includes an enhanced inflammatory profile termed “inflammaging” that correlates with increased vascular permeability, blood-brain barrier (BBB) leakiness, and basal levels of neuroinflammation(Erickson and Banks, 2019). Collectively, these changes increase viral access to the CNS and promote neurotropic infections while also impairing antiviral immune responses(Adesse et al., 2022). CD8^+^ T cells prevent lethality from neurotropic viral infections(Shrestha et al., 2006, 2012; Steinbach et al., 2016; Garber et al., 2019), but with age these responses shrink in number, diversity, and quality(Goplen et al., 2021; Funk et al., 2021). Our previous work showed that aging impaired the recruitment and activation of antiviral CD8^+^ T cells in the CNS during WNV infection, which resulted in loss of virologic control in the CNS and increased lethality in the aged animals(Funk et al., 2021). Furthermore, reports from convalescent COVID-19 patients showed a correlation between long COVID diagnosis and decreased levels of CD8^+^ T cells circulating in peripheral blood(Odak et al., 2020), further suggesting that advanced age results in increased severity of viral infection. However, whether age-associated alterations in the antiviral CD8^+^ T cell response contribute to neurological manifestations of disease or post-infectious cognitive impairment following coronavirus infection remains unclear.

To examine the effect of advanced age on coronavirus-specific CD8^+^ T cell responses, we established a mouse model of respiratory coronavirus infection using MHV-A59. Similar to SARS-CoV-2 MHV-A59 is a member of the subgroup 2a β-coronavirus clade(Graham et al., 2013), but unlike SARS-CoV-2 MHV-A59 is a natural mouse pathogen that readily infects wildtype mice(Körner et al., 2020). Using this model of respiratory coronavirus infection, we found that advanced age significantly increases clinical illness and lethality to MHV-A59, with aged animals harboring increased levels of replicating virus in the CNS during acute infection. This increase in CNS viral burden in aged animals was coupled with increased spatial learning impairment, enhanced microglial activation, and heightened infiltration of activated CD8^+^ T cells into the CNS. However, despite harboring numerically more activated CD8^+^ T cells within the brain, a reduced proportion of CD8^+^ T cells infiltrating the aged brain were specific to MHV, suggesting that aging results in a non-specific recruitment of CD8^+^ T cells into the CNS. Furthermore, aged animals demonstrated increased neuronal death within the hippocampus of infected mice and activated CD8^+^ T cells were shown to induce neuronal death directly in primary cell culture. We attributed this neuronal atrophy to CD8^+^ T cell production of IFN-γ, which was elevated in the aged brain, and correlated with neuronal death in primary neuron cultures. Together these data suggest that aging exacerbates the impact of both direct viral infection and antiviral immune responses within the CNS, which contribute to neuronal dysfunction and cognitive decline.

## Results

### Aged mice are more susceptible to respiratory coronavirus infection

We first examined whether advanced age impacted recovery from respiratory coronavirus infection. Using MHV-A59, 8 wk (adult) or 18 mo old (aged) C57BL/6 mice were infected with 10^4^ plaque-forming units (pfu) via intranasal (i.n) inoculum and their survival and recovery assessed out to 30 days post infection (DPI). While 25% of adult animals succumbed to infection, lethality was significantly increased in aged animals which demonstrated 80% mortality (Figure 1A). Furthermore, adult animals displayed mild weight loss and clinical illness during the acute stages of infection (6-8 DPI) (Figure 1B, C) that was resolved by 10 DPI, at which time animals began to rapidly gain weight and showed no additional signs of clinical illness (Figure 1B-C). Comparatively, MHV-infected aged animals showed significantly increased weight loss compared to either mock infected aged animals or MHV-infected adult animals and did not recover their initial weight by 30 DPI. Additionally, aged animals showed more severe clinical signs of illness during the acute stages of infection with a greater percentage of animals exhibiting clinical scores of 4 and 5 and weight loss persisting out to 12 DPI (Figure 1B-C).

**Figure 1.**
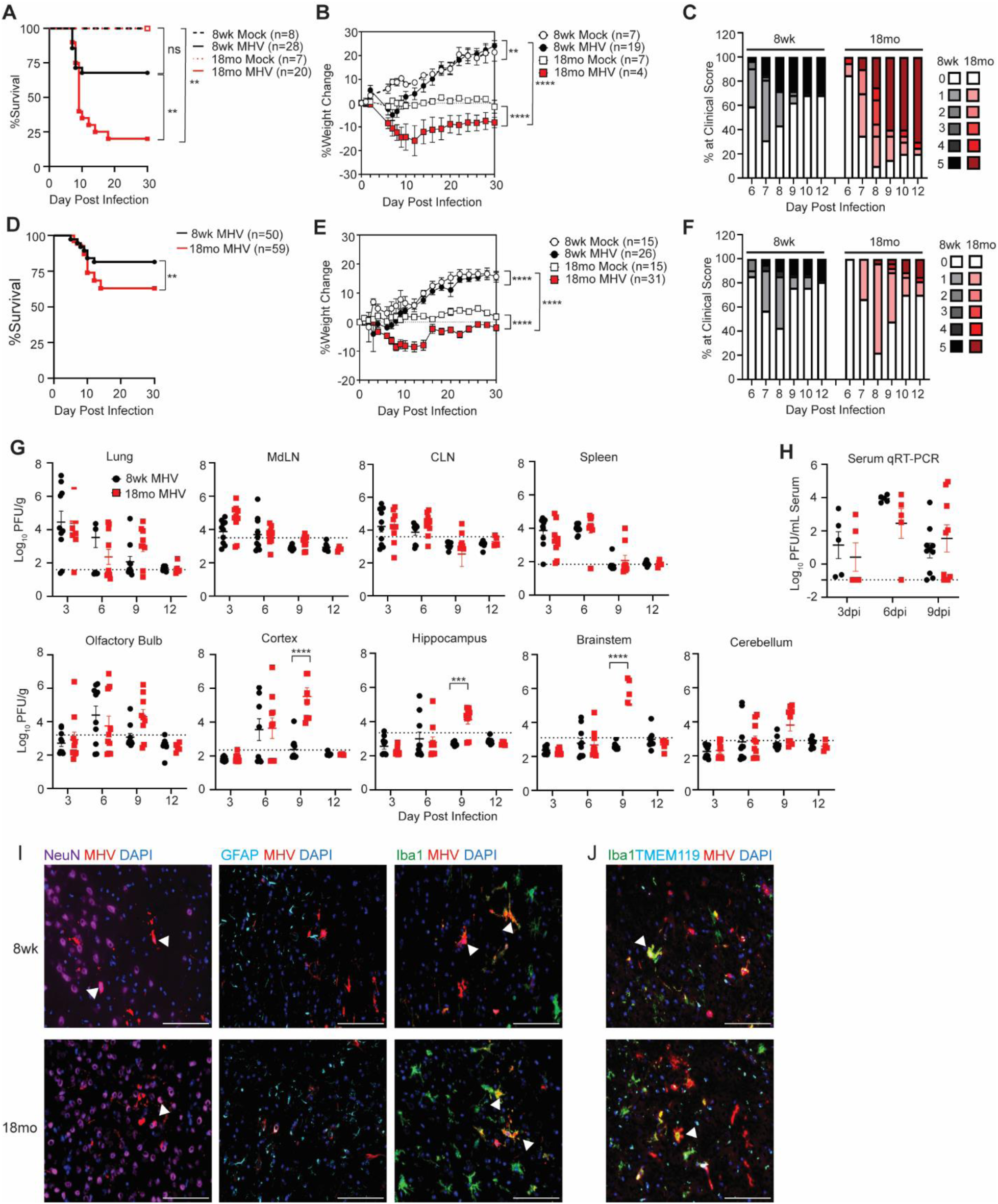
Aged mice are more susceptible to severe respiratory coronavirus infection. 8 wk adult or 18 mo aged C57BL/6 mice were inoculated with 10^4^ pfu (**A-C**) or 10^3^ pfu (**D-J**) MHV-A59 or HBSS (mock infected) i.n. then monitored for mortality (**A, D**), weight change (**B, E**), and clinical illness (**C, F**) for 30 DPI. Clinical scores of adult and aged animals infected with 10^4^ pfu (**C**) or 10^3^ pfu (**F**) MHV-A59 during acute stages of illness were scored as 0= subclinical, 1= hunched/ruffled fur, 2= altered gait/slow movement, 3= no movement, but responsive to stimuli, 4= moribund, 5= dead. Data were pooled from 3 independent experiments with the sample size indicated. (**G**) Viral loads were quantified by plaque assay on the indicated DPI following infection with 10^3^ pfu MHV-A59 i.n. (**H**) Serum viral loads were determined by qRT-PCR. Data were pooled from 3 independent experiments with each data point representing a single animal. (**I**) Representative IHC of MHV N protein (red) co-stained with NeuN (purple), GFAP (cyan), or Iba1 (green) in adult or aged brains at 6 DPI. NeuN^+^MHV^+^ and Iba1^+^MHV^+^ cells indicated by white arrows. (**J**) Representative IHC of MHV-N protein (red) co-stained with Iba1 (green) and TMEM119 (cyan) in adult or aged brains at 6 DPI. Iba1^+^TMEM119^+^MHV^+^ cells indicated by white arrows. Images acquired at 40X magnification. Scale bar = 100 µm. Statistics conducted in GraphPad prism 9.3.1 software. Survival assessed by Log-rank Mantel-Cox assessment, weight change and viral titer assessment conducted according to two-way ANOVA. *, p<0.05; **, p<0.01; ***, p<0.001; ****, p<0.0001

To facilitate our analysis of mechanisms of recovery from infection, we established a model using a lower viral inoculum (10^3^ pfu). Although aged mice still succumbed to infection at a higher rate than adult mice and exhibited increased weight loss and clinical illness, these differences between the adult and aged animals were lessened when inoculated with the lower viral titer (Figure 1D-F). To determine if increased severity in clinical illness observed in aged animals was attributed to elevated viral load, we examined viral titers in the respiratory tract and secondary lymphoid organs following low dose (10^3^ pfu) viral infection. Interestingly, adult and aged animals demonstrated comparable levels of replicating virus in the lung, lung draining mediastinal lymph nodes (MdLN), cervical lymph nodes (CLN), and spleen (Figure 1G), and viral genome in the serum (Figure 1H), suggesting enhanced clinical illness and lethality in aged animals was not attributed to uncontrolled viral replication in these tissues. As coronaviruses are known to have neurotropic potential, we next examined presence of replicating virus within CNS regions of adult and aged animals during acute infection (Figure 1G). Adult and aged animals demonstrated similar kinetics of viral dissemination to the CNS with virus appearing first in the olfactory bulb at 3-6 DPI, then spreading to the cortex and hippocampus by 6 DPI, and then to the hindbrain (cerebellum and brainstem) by 6-9 DPI (Figure 1G). However, aged animals harbored increased CNS viral burden with significantly increased levels of replicating virus detected within the cortex, hippocampus, and brainstem at 9 DPI, a time at which virus had been cleared from the CNS of adult animals (Figure 1G). Despite this, virus was completely cleared from all assessed sites (peripheral and CNS) by 12 DPI, indicating that both adult and aged animals successfully clear viral infection. Together these data suggest that aged versus adult animals are more susceptible to severe outcomes from respiratory MHV infection, in accordance with enhanced viral infiltration into the CNS.

To determine the cellular targets of MHV-A59 in the CNS, we performed IHC analysis on brain tissue sections from mice at 6 DPI, a time point when virus was detected in both adult and aged cortices. Results show that MHV nucleoprotein (N protein) colocalized with NeuN^+^ neurons and Iba1^+^ myeloid cells, but not GFAP^+^ astrocytes, in the prefrontal cortex of both adult and aged mice (Figure 1I). To differentiate between Iba^+^ brain-resident microglia and brain-infiltrating macrophages, we immunostained with microglia specific marker TMEM119. This showed that MHV N^+^Iba1^+^ cells consisted of both TMEM119^+^ microglia as well as TMEM119^neg^ macrophages (Figure 1J), indicating that MHV infects both CNS resident microglia and CNS infiltrating macrophages, as well as neurons. Together these data indicate that respiratory inoculation of MHV results in active CNS infection in both adult and aged animals, validating its use as a model system to study neurotropic effects of respiratory coronavirus infection.

### Respiratory coronavirus infection results in spatial learning defects and cognitive impairment in aged animals

Previous associations between human coronavirus infections and neurologic manifestations(Ellul et al., 2020), along with our detection of MHV in the CNS following murine respiratory infection (Figure 1), led us to hypothesize that respiratory MHV infection may cause cognitive impairment that is exacerbated by advanced age. To test this, mock- or MHV-recovered 8 wk and 18 mo old animals were subjected to open field assessment to measure locomotor function and anxiety, followed by five sequential days of testing in the Barnes maze to assess spatial learning. Open field assessment found no differences between adult or aged animals regardless of infection status in locomotor function measured by animal speed (Figure 2A) or number of line crossings (Figure 2B), or anxiety measured by the total time spent in the center of the arena (Figure 2C). However, when assessed by Barnes maze, post-MHV aged animals were significantly slower to find the target hole compared to either mock infected age-matched controls or adult animals, regardless of infection status (Figure 2D). Area under the curve analysis of adult vs aged animals normalized to their mock controls demonstrated a significantly increased target latency by aged animals (Figure 2E). While all animals improved over the 5 day training period, MHV-infected aged animals were persistently slower to find the target hole, particularly on testing days 2-4 (Figure 2F). Comparatively, adult animals showed no significant difference in spatial learning following infection (Figure 2D-F), suggesting the observed post-infectious cognitive deficits are exclusive to aged animals.

**Figure 2.**
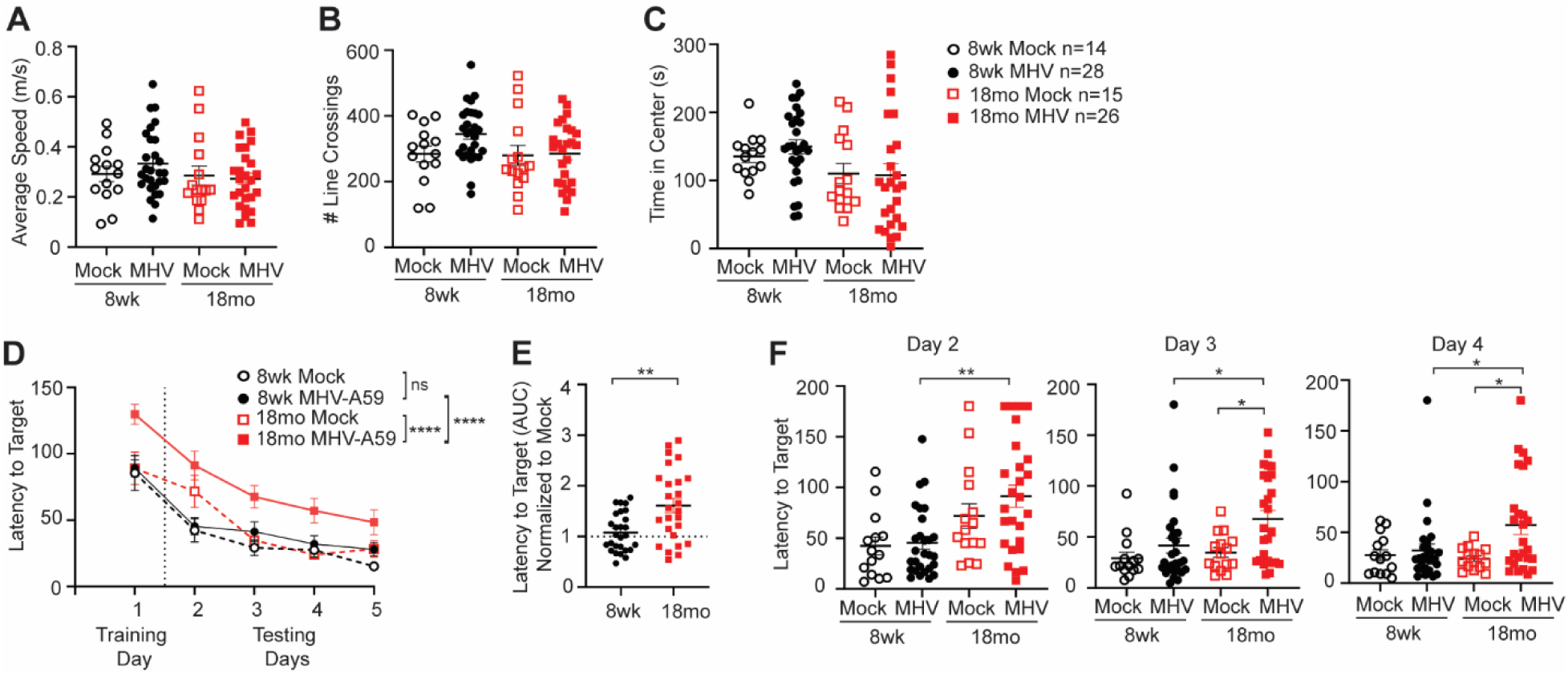
Respiratory coronavirus infection results in spatial learning impairment in aged mice. 8 wk adult or 18 mo aged C57BL/6 mice were inoculated with 10^3^ pfu MHV-A59 or HBSS (mock) i.n., then anxiety and spatial learning were assessed following recovery. **(A-C)** Open field assessment was conducted at 24 DPI and measured the average speed (**A**), total number of lines crossed (**B**) and total time spent in the center region (**C**). (**D-F**) Barnes maze was conducted at 25-29 DPI and latency to target hole was assessed twice each day for 5 days (**D**). (**E**) Overall latency to target was normalized to mock infected age matched controls and area under the curve (AUC) analysis conducted. (**F**) Latency to target by adult vs aged, mock vs infected animals on testing days 2, 3, and 4. Data was pooled from 4 independent experiments with each data point representing a single animal. Statistics according to one-way ANOVA conducted in Graph Pad prism 9.3.1 software. *, p<0.05; **, p<0.01; ***, p<0.001; ****, p<0.0001.

### Aged animals have increased microglial activation following neurotropic MHV infection

Aged microglia typically exhibit a more highly inflammatory or hypersensitive phenotype, resulting in heightened responses to inflammatory stimuli and production of pro-inflammatory cytokines, chemokines, and reactive oxygen species (ROS), which can further enhance the inflammatory microenvironment or have direct neurotoxic effects(Wendimu and Hooks, 2022). Additionally, persistently activated microglia have been shown to promote cognitive dysfunction in both infectious and sterile models of neuroinflammation(Vasek et al., 2016; Di Filippo et al., 2016; Zhang et al., 2021). These pieces of information, coupled with our observation that microglia are a primary target for MHV infection in the CNS (Figure 1I), led to the hypothesis that age-related alterations in the microglial response may contribute to the spatial learning impairment observed in aged animals (Figure 2). To examine the microglial response, we performed flow cytometry analysis on cells isolated from the cortices of adult and aged mice at acute (12 DPI) and post-acute (30 DPI) infection periods. We quantified the total number of CD11b^+^CD45^+^ cells, encompassing both CNS resident microglia and peripheral infiltrating macrophages and monocytes, and the total number of P2RY12^+^ cells, which were specifically microglia. At 12 DPI, we found no difference in the number of CD11b^+^CD45^+^ myeloid cells or P2RY12^+^ microglia between adult or aged animals (Figure 3A, B); however, microglia present in the aged brain showed an increased expression of activation markers MHC II (Figure 3C, D) and CD68 (Figure 3C, E). This heightened microglial activation was sustained at 30 DPI, despite aged animals again having a comparable total number of CD11b^+^CD45^+^ myeloid cells and P2RY12^+^ microglia compared to adult controls (Figure 3F, G). Nonetheless, microglia in the aged brain showed elevated expression of MHC II (Figure 3H, I) and CD68 (Figure 3H, J) compared to those present in adult brains.

**Figure 3.**
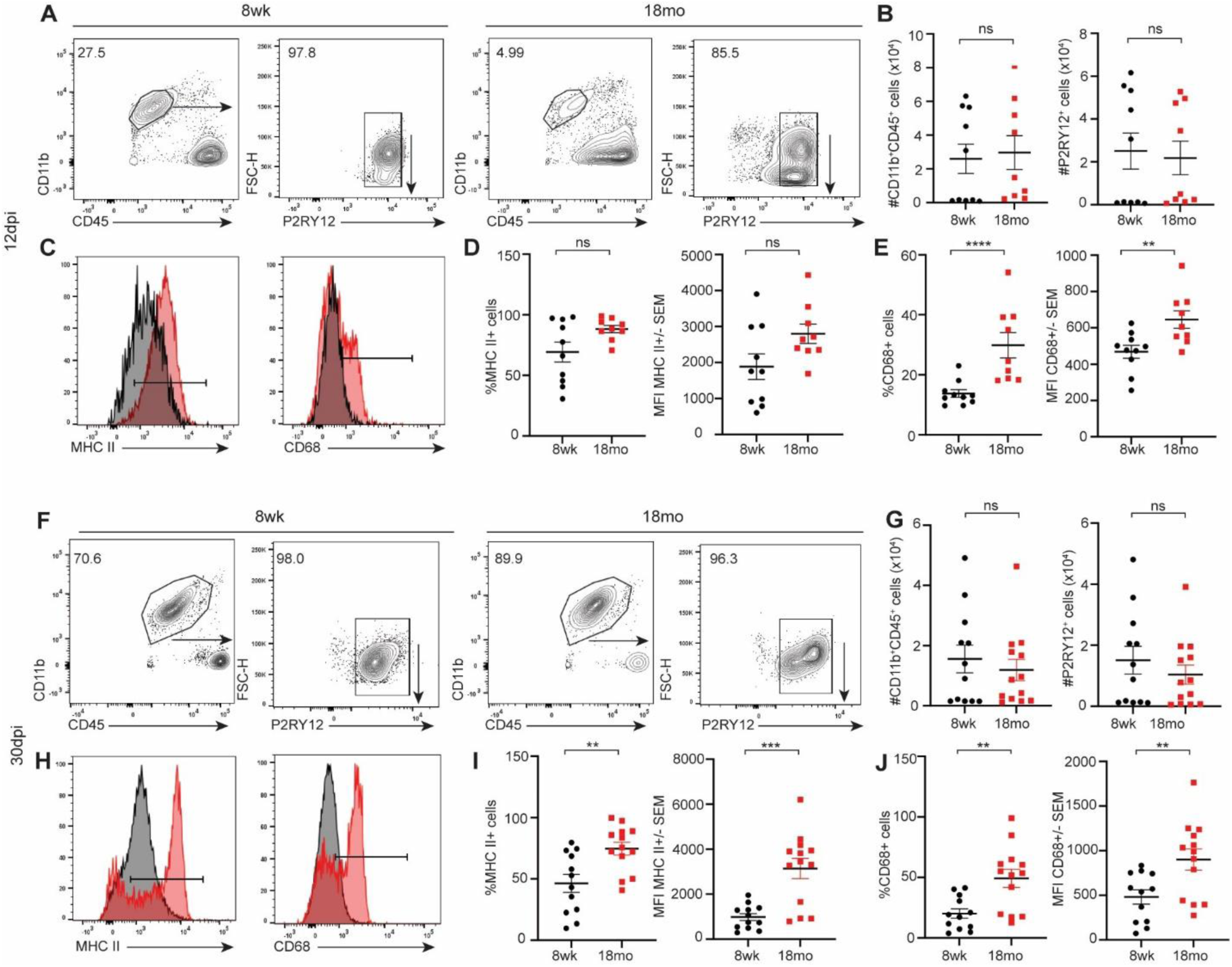
Aged mice have enhanced microglial activation compared to adult mice during respiratory coronavirus infection. 8 wk adult or 18 mo aged C57BL/6 mice were infected with 10^3^ pfu MHV-A59 and microglial activation was assessed at (**A-E**) 12 DPI and (**F-J**) 30 DPI. **(A, F)** Representative flow cytometry of CD11b^+^CD45^+^ microglia and CD11b^+^CD45^+^P2RY12^+^ microglia 12 and 30 DPI. (**B, G**) Quantification of total number of cellular populations. (**C, H**) Histograms of MHC II and CD68 expression by P2RY12^+^ microglia at 12 and 30 DPI. (**D, I**) Quantification of frequency and mean fluorescence intensity (MFI) of MHC II^+^ P2RY12^+^ cells. (**E, J**) Quantification of frequency and MFI of CD68^+^ P2RY12^+^ cells. Data were pooled from 3 independent experiments with each data point representing a single animal. Statistics according to unpaired student t-test conducted in Graph Pad prism 9.3.1 software. *, p<0.05; **, p<0.01; ***, p<0.001; ****, p<0.0001.

Activated microglia target and phagocytose local synapses, disrupting synaptic connections and resulting in cognitive impairment in the post-infectious brain(Vasek et al., 2016). To determine if heightened microglial activation observed in aged animals previously infected with MHV resulted in synapse elimination, we examined expression of the presynaptic marker, Synaptophysin, in the hippocampus of adult and aged, mock or MHV infected mice at 30 DPI. We saw no difference in the percent area of Synaptophysin staining, and there was no colocalization of Iba1^+^ cells with Synaptophysin^+^ synapses in the aged brain after infection, as determined by Pearson Correlation Coefficient (PCC; Supplemental Figure 1). These results suggest that microglia-mediated synaptic loss was not a significant contributor to cognitive decline in aged animals in this model.

We also examined demyelination as a potential cause of spatial learning impairment, as MHV has been shown to result in demyelinating disease when delivered by direct intracranial inoculation(Lavi et al., 1984). We used Luxol fast blue staining and examined corpus callosum width at three different regions (caudal, medial, and rostral) as a measure of myelin density, and found no difference in corpus callosum size regardless of age or treatment group (Supplemental Figure 2). Furthermore, we found no difference in the proportion of myelin basic protein (MBP) in the corpus callosum (Supplemental Figure 2), suggesting that while MHV can result in demyelination, we do not observe myelin loss in our model system and that an alternative mechanism contributes to the spatial learning impairment observed in aged animals.

### Aged mice have enhanced CD8^+^ T cell responses compared to adult animals following recovery from respiratory coronavirus infection

Increased severity of neurotropic viral infection and CNS viral burden has been attributed to deficiencies in the aged antiviral CD8^+^ T cell response(Funk et al., 2021). Furthermore, brain infiltrating CD8^+^ T cells contribute to cognitive decline via direct and indirect mechanisms(Reagin and Funk, 2022). To examine the CD8^+^ T cell response within the brain during acute (12 DPI) and recovery (30 DPI) stages of MHV infection, we performed flow cytometry analysis on cortical tissue collected from adult and aged animals. Interestingly, we found a significant increase in the infiltration of both CD4^+^ and CD8^+^ T cells into the aged brain compared to adult controls (Figure 4A). The frequency and total number of CD8^+^ T cells was significantly elevated in the brains of aged animals compared to adult controls at both acute (Figure 4B) and recovery (Figure 4C) time points. This was unexpected as aging is conventionally associated with decreased cellular immunity to viral infection(Funk et al., 2021); however, despite harboring increased numbers of CD8^+^ T cells, a significantly lower percentage of CD8^+^ T cells within the aged brain were MHV-specific (Figure 4D). We examined MHV antigen specificity by brain infiltrating CD8^+^ T cells to the immunodominant MHV S protein using an MHC I tetramer, and found that adult animals harbored increased antigen-specific MHV S-tet^+^ cells both acutely (11% vs 4%, Figure 4E), as well as following recovery from infection (10% vs 2%, Figure 4F). However, due to the increase in the total number of brain-infiltrating CD8^+^ T cells, aged animals demonstrated no difference in the total number of MHV-specific CD8^+^ T cells in the brain acutely at either time point (Figure 4E, F).

**Figure 4.**
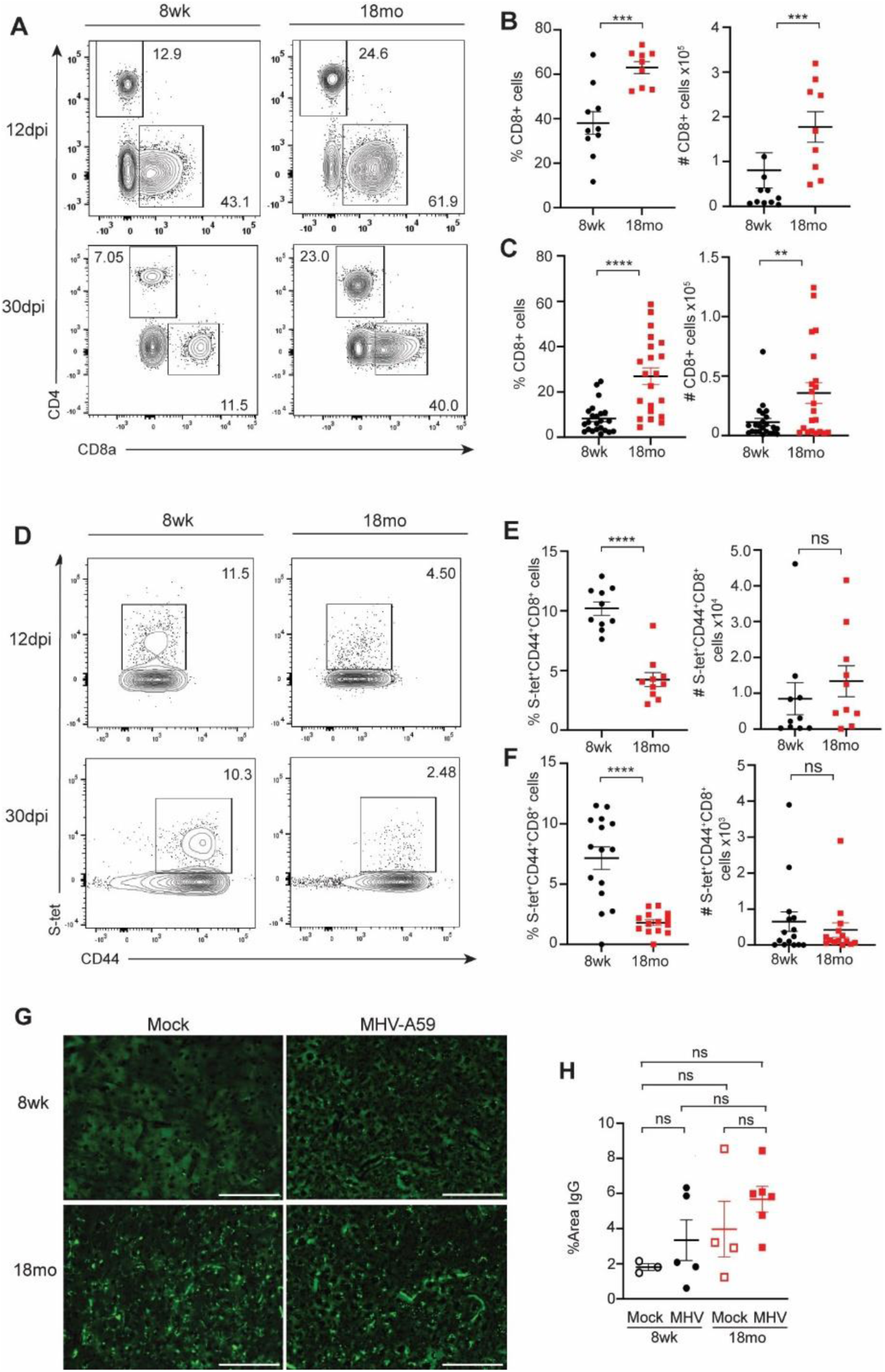
Aged mice have enhanced infiltration of CD8^+^ T cells into the CNS following respiratory coronavirus infection. 8 wk adult or 18 mo aged C57BL/6 mice were infected i.n. with 10^3^ pfu MHV-A59 and CD4^+^ and CD8^+^ T cell response were assessed in the cortex at 12 and 30 DPI. (**A)** Representative flow cytometry plots of CD4^+^ and CD8^+^ cells in the brain 12 DPI (**top**) and 30 DPI (**bottom**). (**B, C**) Quantification of the frequency and total number of CD8^+^ T cells at each time point. (**D)** Representative flow cytometry plots of CD44 and MHV S-tetramer on CD8^+^ T cells from the cortices of adult or aged animals at 12 DPI (**top**) and 30 DPI (**bottom**). (**E, F)** Quantification of frequency and total number of CD44^hi^S-tet^+^ cells. Data were pooled from 3 or 4 independent experiments with each data point representing an individual animal. (**G**) Representative IHC staining for IgG in the cortex of 8 wk adult or 18 mo, mock or MHV-A59 infected animals. Images taken at 40X magnification. Scale bar = 100 µm. (**H**) Quantification of total percent area of IgG in the CX. Data representative of 1 experiment with each data point representing a single animal. Statistics according to unpaired Student T-test (flow cytometry) or one-way-ANOVA (IHC) conducted in Graph Pad prism 9.3.1 software. *, p<0.05; **, p<0.01; ***, p<0.001; ****, p<0.0001.

This enhanced CD8^+^ T cell response observed in aged animals was not restricted to theCNS, as aged animals harbored increased activated CD44^hi^CD8^+^ T cells in all sites assessed, including the lung, draining MdLN, CLN, and spleen at both low and high viral inoculum (Supplemental Figure 3A, B). This is likely not driven by enhanced viral burden, as equivalent levels of replicating virus were detected in those peripheral sites at all time points assessed (Figure 1G). Although actively replicating virus was undetectable in both adult and aged animals by 12 DPI, we tested whether viral genome persisted beyond the 12 DPI time point, which may drive continual activation and recruitment of CD44^hi^CD8^+^ T cells(Schøller et al., 2019). Assessment of viral genome by qRT-PCR detected comparable levels of MHV in the lung, cortex, and hippocampus of adult and aged animals at 12 DPI, indicating the presence of either non-replicating virus or replicating virus that was below the limit of detection of the plaque assays (Figure 1). However, no viral genome was detected in adult or aged animals by 30 DPI (Supplemental Figure 3C), suggesting any persisting viral reservoirs are cleared by this time. Together, these data suggest that persistent viral infection was likely not responsible for the increased CD8^+^ T cell response observed in the brain of aged animals. However, we did observe enhanced disruption of the BBB in aged animals following MHV infection compared to mock infected controls (Figure 4G, 4H), albeit not to statistically significant levels. This suggests that enhanced barrier disruption associated with advanced age may permit the entry of non-specific CD8^+^ T cells into the CNS.

### Respiratory MHV infection results in neuronal death

Infiltration of CD8^+^ T cells into the brain can contribute to spatial learning impairment directly via apoptosis of infected neurons(Reagin and Funk, 2022). CNS infiltrating CD8^+^ T cells mediate clearance of viral reservoirs by promoting neuronal apoptosis directly via Fas-ligand interactions(Shrestha and Diamond, 2007), as well as through signaling via TNF-related apoptosis-inducing ligand (TRAIL)(Shrestha et al., 2012). Additionally, secretion of cytotoxic granules including perforin(Shrestha et al., 2006) and granzyme(Wu et al., 2021) can result in neuronal atrophy via activation of caspase-3. Therefore, to determine if the spatial learning impairment observed in aged animals correlated with neuronal apoptosis, we measured TUNEL^+^ neurons within regions of the hippocampus that are responsible for spatial learning and memory: the dentate gyrus (DG), CA1, and CA3. We examined these regions in adult and aged animals acutely and post recovery from MHV infection. We found TUNEL^+^NeuN^+^ cells in both the DG and CA3 regions of the hippocampus in both adult and aged animals (Figure 5A, B), but no TUNEL staining was detected in the CA1 (data not shown). More TUNEL^+^ cells were observed at the acute timepoint (12 DPI) versus the recovery timepoint (30 DPI) in both age groups, suggesting neuronal death occurs with acute infection and likely resolves with recovery and viral clearance. Results showed a greater percent area positive for TUNEL in aged versus adult DG and CA3 at 12 DPI, and to a lesser extent at 30 DPI (Figure 5C, D). Using PCC and Mander’s overlap coefficient, we saw significantly greater colocalization between TUNEL and NeuN in aged animals in the CA3 at all timepoints and in the DG at most timepoints (Figure 5E, F). Together, these data indicate increased neuronal death in aged versus adult animals, particularly during acute infection. This increased neuronal death observed in aged animals was accompanied by a decrease in neuronal regeneration, as aged animals showed a reduction in neurogenesis compared to adult controls (as assessed by doublecortin (DCX) staining), regardless of infection status (Supplemental Figure 4). This suggests that MHV-A59 infection results in neuronal atrophy within the hippocampus of both adult and aged animals acutely; however, this is more severe and sustained in aged animals.

**Figure 5.**
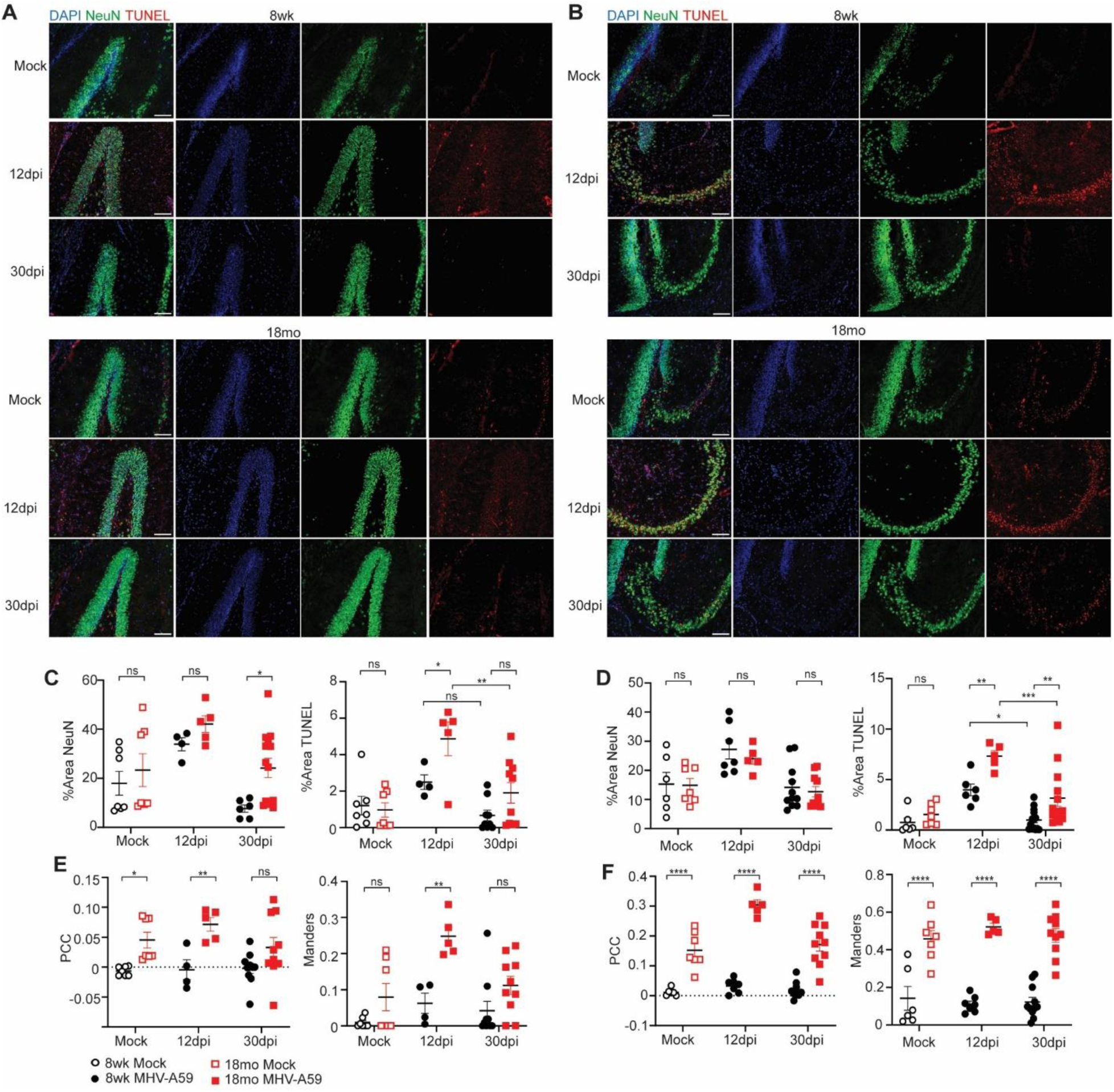
MHV-A59 respiratory infection results in enhanced neuronal death in aged hippocampus. Representative IHC of DAPI, NeuN, and TUNEL in the (**A**) dentate gyrus and (**B**) CA3 of 8 wk or 18 mo old animals at 12 and 30 DPI compared to mock infected controls. Images taken at 20X magnification. Scale bar = 100 µm. (**C, D**) Quantification of total percent area of NeuN and TUNEL in the (**C**) DG or (**D**) CA3. Colocalization between NeuN and TUNEL measured by PCC and Mander’s overlap coefficient in the (**E**) DG and (**F**) CA3. Data were pooled from 2 independent experiments with each data point representing a single animal. Statistics according to unpaired one-way ANOVA conducted in Graph Pad prism 9.3.1 software. *, p<0.05; **, p<0.01; ***, p<0.001; ****, p<0.0001.

### Activated CD8^+^ T cells Mediate Neuronal Death In Vitro

To determine if CD8^+^ T cells directly contributed to the neuronal death observed in aged animals, we first examined localization of brain-infiltrating CD8^+^ T cells. We conducted IHC analysis of adult and aged brains at 12 and 30 DPI and easily identified CD8^+^ T cells at 12 DPI in the cortex and DG in both adult and aged brains (Figure 6A). However, by 30 DPI, CD8^+^ T cells were absent from the hippocampus entirely and were only found in the cortex of adult and aged animals (Figure 6B). This was interesting, as no CD8^+^ T cells were identified within the hippocampal CA3 region, where the majority of neuronal apoptosis was observed (Figure 5D). This suggested that an indirect soluble factor may be the primary mediator of neuronal death, rather than direct mechanisms of CD8^+^ T cell directed neuronal killing. We next examined cytokine production and cytolytic function by CD8^+^ T cells isolated from the adult or aged brain of MHV-infected or mock infected controls following recovery from infection (30 DPI). We found MHV-infected adult mice had a higher proportion of IFN-γ (Figure 6C, 6D) and TNF (Supplemental Figure 5A, D) producing cells compared to their mock infected counterparts. However, the proportion of IFN-γ producing cells was elevated in aged animals even at basal levels, and increased moderately with infection (Figure 6C, D), suggesting CNS infiltrating CD8^+^ T cells in aged animals had a more highly activated phenotype even at baseline. However, there was no significant increase in TNF production in aged animals (Supplemental Figure 5D). Both adult and aged animals showed a moderate increase in granzyme B (GZMB) production following infection, but this was not statistically significant (Supplemental Figure 5B, E). Additionally, there was no observed differences in production of perforin between adult or aged animals irrespective of infection status (Supplemental Figure 5C, F). Together, this suggested elevated production of IFN-γ by CD8^+^ T cells may contribute to neuronal atrophy within the CNS following infection.

**Figure 6.**
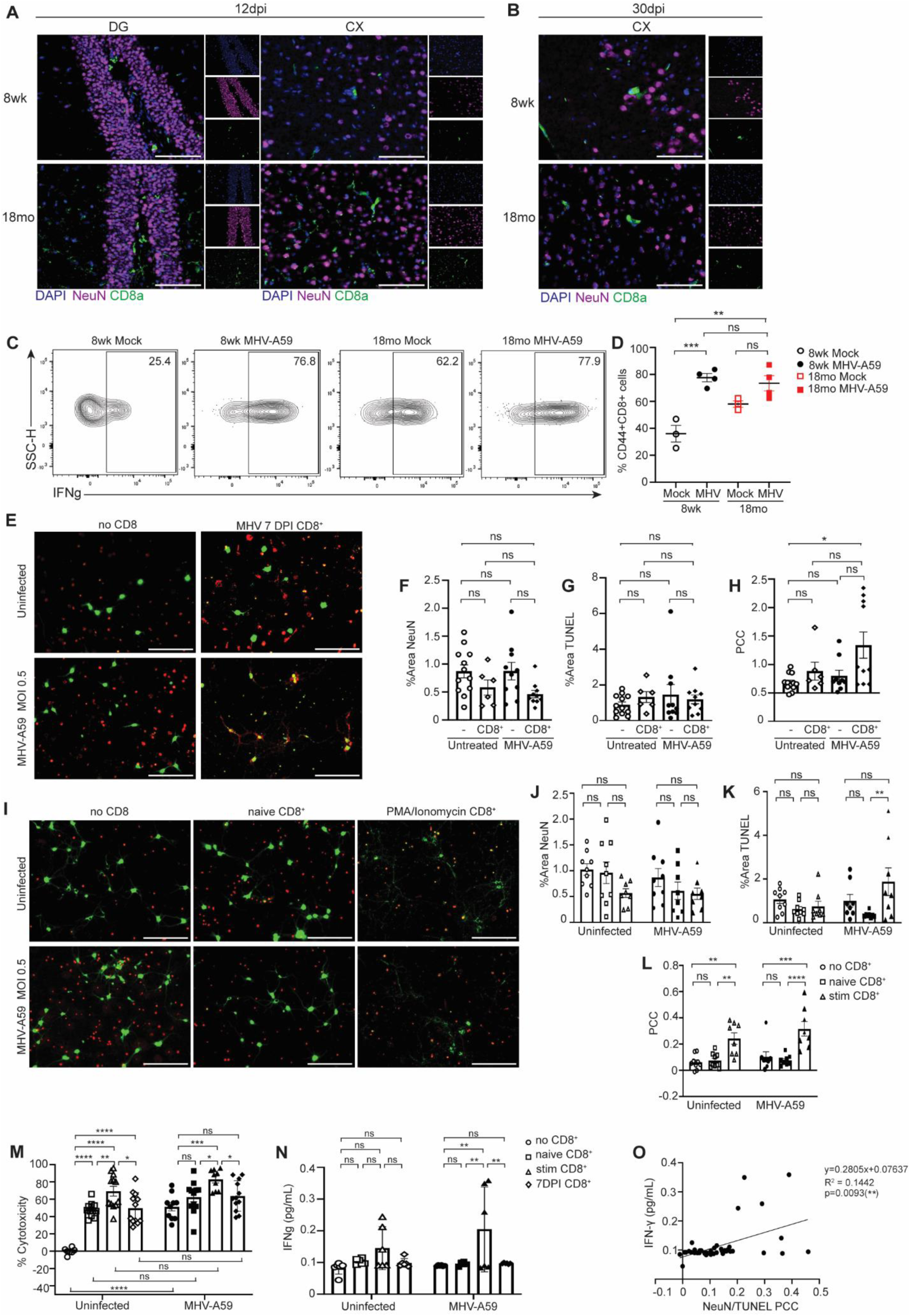
IFN-γ produced by CD8^+^ T cells mediates neuronal apoptosis following MHV-A59 infection. (**A**) Representative IHC of DAPI, NeuN, and CD8a in the DG and cortex (CX) of 8 wk or 18 mo old animals at (**A**) 12 DPI (**B**) and 30 DPI. (**B**) Representative flow cytometry plots of IFNγ^+^ CD8^+^ T cells isolated from the brains of 8 wk or 18 mo old, mock or MHV-A59 infected animals at 30 DPI following 4 hr PMA/ionomycin stimulation. (**C**) Representative flow cytometry and **(D)** quantification of percent CD8^+^CD44^+^ cells positive for IFN-γ. (**E**) Representative immunocytochemical (ICC) staining for NeuN and TUNEL of primary cortical neurons following no treatment or infection with MHV-A59 MOI 0.5, with or without co-culture with CD8^+^ T cells purified from the spleen of an 8 wk old animal at 7 DPI with 10^3^ pfu MHV-A59. (**F-H**) Quantification of ICC images: (**F**) percent area NeuN, (**G**) percent area TUNEL, (**H**) colocalization between NeuN and TUNEL using PCC. (**I**) Representative ICC of NeuN and TUNEL staining of primary embryonic cortical neurons following no treatment or infection with MHV-A59 MOI 0.5, with or without naïve or PMA/ionomycin stimulated CD8^+^ T cells purified from the spleen of an 8 wk old uninfected animal. (**J-L**) Quantification of ICC images: (**J**) percent area NeuN, (**K**) percent area TUNEL, and (**L**) colocalization between NeuN and TUNEL using PCC. (**M**) Percent cytotoxicity of primary cortical neurons following no treatment or infection with MHV-A59 MOI 0.5, with or without naïve or PMA/ionomycin stimulated CD8^+^ T cells purified from the spleen of an 8 wk uninfected animal, or CD8^+^ T cells purified from the spleen of an 8 wk MHV-A59 infected animal 7 DPI. (**N**) Total concentration of IFN-γ (pg/mL) assessed by ELISA from co-culture supernatant collected from panel M. (**O**) Linear regression analysis of IFN-γ concentration and TUNEL/NeuN PCC. Data are representative of three independent experiments with each data point representing an individual sample. All images were taken at 40X magnification and scale bar = 100 µm. 3 images were captured per sample. Statistics according to unpaired one-way ANOVA conducted in Graph Pad prism 9.3.1 software. *, p<0.05; **, p<0.01; ***, p<0.001; ****, p<0.0001.

To further assess the ability of CD8^+^ T cells to contribute to neuronal death, we utilized an *in vitro* co-culture system in which CD8^+^ T cells were isolated from the spleens of adult animals at 7 DPI, then co-cultured with naïve embryonic cortical neurons. Neurons were either untreated or infected with MHV at multiplicity of infection (MOI) 0.5, then purified CD8^+^ T cells were added at a 1:1 ratio 24 hrs later. Neurons and T cells were incubated for an additional 24 hrs, at which point neurons were assessed for apoptosis via TUNEL staining. We found that infection with MHV alone did not cause neuronal death, as there was no change in the total percent area of NeuN^+^ neurons (Figure 6E, 6F), and we detected minimal TUNEL staining observed in either uninfected or MHV-infected neurons (Figure 6E, G). However, co-culturing infected neurons with 7 DPI CD8^+^ T cells resulted in an increase in the colocalization of TUNEL^+^ cells with NeuN^+^ neurons, suggesting neuronal apoptosis (Figure 6E, 6H). Interestingly, this CD8^+^ T cell mediated neuronal killing was not antigen-specific. CD8^+^ T cells isolated from an uninfected animal, which were stimulated with PMA/ionomycin to induce high levels of non-specific activation, caused neuronal death *in vitro* while uninfected CD8^+^ T cells left unstimulated did not (Figure 6I). In fact, infection with MHV or treatment with uninfected/unstimulated CD8^+^ T cells resulted in no difference in the proportion of NeuN^+^ cells (Figure 6J) or TUNEL^+^ cells (Figure 6K); however, addition of PMA/ionomycin-stimulated CD8^+^ T cells to either infected or uninfected neurons resulted in a significant increase in colocalization between NeuN^+^ and TUNEL^+^ cells, which was not observed following the addition of uninfected/unstimulated CD8^+^ T cells (Figure 6L). These data suggested that activated CD8^+^ T cells caused neuronal apoptosis, independent of CD8^+^ T cell antigen specificity or neuronal cell infection. As a more holistic measure of cell death, we assessed CD8^+^ T cell induced neuronal death in our co-culture system using a high-throughput fluorometric cytotoxicity assay. Primary neuron cultures were mock- or MHV-infected at MOI 0.5, then 24 hrs later neurons were co-cultured with either naïve, PMA/ionomycin-stimulated, 7 DPI, or no CD8^+^ T cells. In contrast to the TUNEL staining, this fluorometric assay detected reduced viability in MHV-infected compared to uninfected neurons in the absence of co-cultured T cells (Figure 6M). This could be attributed to the increased sensitivity of the fluorometric assay to early cell damage or non-apoptotic mechanisms of cell death, which may be undetected by TUNEL stain(Kari et al., 2022). We also found reduced viability in uninfected neuronal co-cultures containing naïve, PMA/ionomycin stimulated, or MHV 7 DPI T cells (Figure 6M). While all three CD8^+^ T cell treatment groups elicited neuronal death *in vitro* compared to untreated controls, PMA/ionomycin stimulated CD8^+^ T cells elicited the highest level of neuronal death, regardless of infection status. To further investigate the role of IFN-γ in neuronal cell death, we quantified the levels of IFN-γ in this system. Results showed that neuron co-cultures containing PMA/ionomycin stimulated CD8^+^ T cells expressed the highest levels of secreted IFN-γ, as assessed by ELISA (Figure 6N), and IFN-γ levels positively correlated with colocalization of TUNEL/NeuN (Figure 6O), suggesting that CD8^+^ T cell production of IFN-γ contributes to neuronal apoptosis. Together, these data demonstrate that enhanced recruitment of activated CD8^+^ T cells into the aged CNS following neurotropic coronavirus infection causes neuronal atrophy, which correlates with spatial learning deficits and post-infectious cognitive decline observed in aged individuals.

## Discussion

The pandemic caused by respiratory coronavirus SARS-CoV-2 is entering its fourth year, and much remains unknown regarding the cellular mechanisms by which coronavirus infection impacts cognitive health. Emerging studies have reported that approximately 1 in 3 patients who have recovered from infection will experience persistent symptoms of long COVID, including cognitive complications(Taquet et al., 2021). While long COVID is typically associated with increased severity of acute disease, its presence in mild and asymptomatic patients suggests that long COVID diagnoses may be under-reported(Ma et al., 2023). The incidence of long COVID disproportionately affects individuals of advanced age(Taquet et al., 2021), putting this population at a higher risk for not only severe infection but also persistent post-infectious symptoms. With aged individuals comprising approximately 20% of the world’s population in the coming years(United Nations, 2023), it is critical that we understand mechanisms through which advanced age impacts the lasting effects of coronavirus infection in order to develop therapeutic intervention strategies.

Using a murine model of coronavirus (MHV-A59), this study demonstrated that respiratory coronavirus infection resulted in enhanced mortality and viral dissemination to the CNS in aged versus adult mice. Furthermore, our data revealed that respiratory coronavirus infection resulted in spatial learning impairment, which correlated with increased infiltration of antiviral CD8^+^ T cells to the brain, suggesting a role in post infectious cognitive decline. Consistent with patient reports of exacerbated cognitive decline in aged individuals(Sullivan and Fischer, 2021), aged mice showed significant increase in latency to find the target hole. In agreement with previous publications, aged animals in this study showed delayed spatial learning even in the absence of infection(Yang et al., 2019), suggesting that aging-intrinsic factors contribute to spatial learning deficits that are likely exacerbated with infection. This study used exclusively male mice because early patient reports pointed to aged men as the most susceptible to severe viral infection(Bwire, 2020). More recent reports indicate that women are at greater risk of developing long COVID, possibly due to their enhanced immune response(Bai et al., 2022), which will be an interesting line of future inquiry.

Previous studies have shown that MHV-A59 causes significant demyelinating disease when delivered by direct intracranial inoculation(Lavi et al., 1984), an effect that is greatly amplified when substituted with the highly neurovirulent MHV-JHM strain (Lavi and Constantinescu, 2005). However, we observed no signs of demyelination in this model of intranasal (i.n.) inoculation, as measured by corpus collosum width with Luxol fast blue or immunostaining with MBP, suggesting demyelination was not a significant contributor to the cognitive deficits observed in this study. We examined other molecular mechanisms which have been shown to drive cognitive decline in other model systems including microglia-mediated synaptic elimination(Vasek et al., 2016) and direct neuronal apoptosis(Moujalled et al., 2021), both of which have been shown to increase concurrent with advanced age and inflammation(Wendimu and Hooks, 2022). We found little to no difference in synaptic staining in adult or aged animals regardless of infection status, but a significant amount of neuronal death specifically within the hippocampal CA3 and DG regions following MHV infection. Uninfected adult animals showed no signs of neuronal death, but this increased following acute MHV infection then dissipated following recovery from infection. Conversely, aged animals demonstrated significant neuronal apoptosis within the hippocampus even in the absence of infection; however, this increased with acute infection and was sustained following recovery. This apoptosis was coupled with a decrease in neuronal regeneration in aged but not adult animals. These data suggest that neuronal apoptosis within the hippocampal trisynaptic circuit, which is responsible for visuospatial memory formation(Basu and Siegelbaum, 2015), in tandem with a lack of regeneration, drives spatial learning deficits observed in aged animals post-infection. While we observed neuronal death in adult animals acutely, they showed neuronal regeneration during the recovery phase, potentially restoring the hippocampal trisynaptic circuit and ameliorating perturbations in visuospatial memory.

Our data point to CNS infiltrating inflammatory CD8^+^ T cells as drivers of neuronal apoptosis observed in the hippocampus. Under homeostatic conditions, CD8^+^ T cells are largely restricted from the CNS; however, with infection or inflammation CNS barriers become leaky, and coupled with production of inflammatory mediators by resident glial cell populations, CD8^+^ T cells are recruited to the CNS perivascular space and parenchyma(Smolders et al., 2013). During infection in the CNS, CD8^+^ T cells are critical for restricting viral replication(Shrestha et al., 2006; Shrestha and Diamond, 2007) and recrudescence(Khanna et al., 2003), via production of perforin and granzyme, as well as cytokines IFN-γ and TNF. However, increasing evidence implicates activated inflammatory CD8^+^ T cells in cognitive disease(Reagin and Funk, 2022). Models of neurotropic flavivirus infection have shown CNS infiltrating CD8^+^ T cells as both protective against neurotropic infection and harmful to glial and neuronal health(Shrestha and Diamond, 2004). CD8^+^ T cells specific to either Zika virus or WNV were shown to directly contribute to post-infectious cognitive decline via their production of IFN-γ, which enhanced microglial mediated synaptic elimination(Garber et al., 2019). Here, we demonstrated that infiltration of activated CD8^+^ T cells into the CNS following neurotropic coronavirus infection correlated with post-infectious cognitive impairment that was more severe in advanced age.

The CNS of aged animals demonstrated enhanced infiltration of both CD4^+^ and CD8^+^ T cells both acutely and following recovery from infection, as well as increased CD44^hi^CD8^+^ T cells in all peripheral sites assessed, including the lungs and secondary lymphoid organs. These results were contradictory to our previous study using WNV, which showed a reduced antiviral CD8^+^ T cell response within the CNS of aged mice(Funk et al., 2021). Although we did not directly compare WNV and MHV in this study, we hypothesize that these distinct responses may be due to differences in viral neurotropism and antiviral immunity. Whereas once in the CNS, WNV specifically infects neurons(Guarner et al., 2004; Omalu et al., 2003), we saw that MHV infected myeloid cells, both brain-resident microglia and infiltrating macrophages, as well as neurons. We posit that this ability to productively infect myeloid cells may promote relatively greater inflammatory signaling in MHV infected mice, thus enhancing the peripheral CD8^+^ T cell response and recruiting greater numbers of CD8^+^ T cells to the brain. In support of this, previous studies investigating the peripheral immune response to MHV and WNV have shown distinct inflammatory cytokine expression. Whereas i.n. inoculation of MHV into aged mice showed a hyperinflammatory response indicated by increased serum levels of TNF, IL-1β, and IL-6(Ryu et al., 2021), subcutaneous inoculation of WNV showed decreased levels of inflammatory cytokines in the draining lymph nodes of aged versus adult mice(Richner et al., 2015). This suggests a global alteration in the aged antiviral response to MHV versus WNV infection, but the underlying mechanism driving these divergent responses remains unknown.

Despite the greater numbers of infiltrating CD8^+^ T cells, the majority of the CNS infiltrating CD8^+^ T cells in the aged brain were not antigen-specific to the immunodominant MHV S-protein. Considering that aging is associated with an increase in low grade systemic inflammation even under homeostatic conditions(Goplen et al., 2021), it is likely that the enhanced inflammatory microenvironment within the aged brain, coupled with enhanced BBB leakiness observed with aging, could be driving non-specific recruitment of CD8^+^ T cells into the CNS following infection. Our data using an *in vitro* co-culture experimental design demonstrated that activated CD8^+^ T cells can induce neuronal death even in the absence of a viral cognate antigen. While CD8^+^ T cells isolated from an MHV infected mouse at 7 DPI were capable of inducing apoptosis of MHV infected neurons, CD8^+^ T cells isolated from an uninfected mouse, when stimulated with PMA/ionomycin, also caused neuronal death to both MHV infected and uninfected primary neurons. This suggests that activated CD8^+^ T cells within the CNS can induce neuronal apoptosis of both infected target cells and uninfected bystander cells. CD8^+^ T cell mediated neuronal targeting is typically restricted through T cell intrinsic mechanisms including expression of inhibitory receptors(Suvas et al., 2006), or through niche specific environmental factors including glial cell expression of inhibitory ligands such as PD-L1(Manenti et al., 2022), and neuronal downregulation of MHC-I expression(Miller et al., 2016), but how these processes change with infection and advanced age are unclear. Thus, considering that we saw large numbers of infiltrating CD8^+^ T cells that were not specific to MHV, and the ability of non-specific CD8^+^ T cells to kill uninfected neurons *in vitro*, these data indicate that non-antigen specific mechanisms likely underlie the CD8^+^ T cell mediated neuronal apoptosis in this system.

We have identified IFN-γ as a key mediator of neuronal death caused by CD8^+^ T cells. We observed heightened levels of IFN-γ in neuron co-cultures containing activated, but not naïve, CD8^+^ T cells which correlated directly with TUNEL^+^ neuronal death. As expected, examination of IFN-γ production by CNS infiltrating CD8^+^ T cells demonstrated that adult CD8^+^ T cells produce copious amounts of IFN-γ following infection compared to mock infected controls; however, CD8^+^ T cells within the brain of aged animals showed elevated levels of IFN-γ production irrespective of infection status. Aged CD8^+^ T cells have a lower threshold of T cell receptor engagement requisite for successful activation, suggesting that aged CD8^+^ T cells maintain a basal level of heightened responsiveness to inflammatory signals(Goronzy et al., 2012). This, coupled with the increased basal levels of inflammation that occur with age, suggests aged CD8^+^ T cells may be poised in a constant inflammatory state which is harmful to neurons upon CNS entry(Goplen et al., 2021). Adult and aged CD8^+^ T cells also displayed increased expression of TNF and GZMB following MHV infection, albeit not to statistically significant levels; therefore, it is likely that IFN-γ alone is not the only contributor to neuronal death in our model system. However, previous studies have shown the impact of IFN-γ on neurons, including regulating neurotransmission(Janach et al., 2022, 2020), neurotoxicity(Mizuno et al., 2008), and neurogenesis(Li et al., 2010). Future experiments will be required to determine the direct mechanisms of CD8^+^ T cell induced neuronal apoptosis in this system and how those may be altered or exacerbated with advanced age.

Bidirectional communication between microglia and CD8^+^ T cells together drive the inflammatory state within the CNS and collectively contribute to post-infectious cognitive decline. Microglia activate infiltrating CD8^+^ T cells, and CD8^+^ T cells cyclically enhance inflammatory cytokine production by microglia. Thus, altered CD8^+^ T cell/microglia interactions likely also contribute to spatial learning impairment and neuronal death in aged animals. Aged animals demonstrated enhanced activation of microglia at both acute and recovery timepoints. While it is possible that this sustained activation could be due to persistent viral infection, no detectable levels of replicating virus were identified in the CNS at 12 DPI, and viral genome was undetectable within the CNS at 30 DPI, suggesting this sustained microglial activation could be inherent to aging itself. Aged microglia exhibit a hypersensitive phenotype, resulting in heightened responses to inflammatory stimuli and production of pro-inflammatory cytokines that can further enhance the inflammatory microenvironment or have direct neurotoxic effects(Wendimu and Hooks, 2022). This expression of inflammatory cytokines also drives infiltration of immune cells, including CD8^+^ T cells, into the CNS(Zhang et al., 2022; Funk and Klein, 2019). Furthermore, once present in the CNS, CD8^+^ T cells are activated by microglia, which cross-present phagocytosed viral antigen, further driving CD8^+^ T cell activation and effector function(Moseman et al., 2020). Therefore, the heightened microglial activation observed in aged animals may amplify the activated CD8^+^ T cells present within the CNS. However, as microglia represent one of the primary CNS resident cell types infected with MHV, the role of reciprocal micgolial-CD8^+^ T cell interactions in contributing to spatial learning impairment in aged animals remains unexplored. Future experiments will seek to uncouple the microglial-CD8^+^ T cell response during viral infection to better understand the bidirectional relationship between aged microglia and antiviral CD8^+^ T cell responses within the aged brain.

The data presented here establish a role for CD8^+^ T cells in post-infectious cognitive decline following respiratory coronavirus infection. These data demonstrate alterations in the antiviral CD8^+^ T cell response in aged animals, which may underlie the severity of coronavirus infection observed in elderly patients. In conclusion, we show that alterations in the aged immune response to MHV-A59 coronavirus infection may contribute to cognitive disease, and may inform improved therapeutic interventions to SARS-CoV-2 and other coronavirus infections.

## Methods

### Virus Preparation

MHV-A59 (VR-764) stocks and producer cell line NCTC-1469 (CCL-9.1) were obtained from the American Type Culture Collection, ATCC (Manassas, VA, USA). NCTC-1469 were maintained in Dulbecco’s modified Eagle’s Medium (DMEM; Life Technologies, cat#11965126, Grand Island, NY) supplemented with 10% heat inactivated horse serum (VWR, cat#103219-558, Suwanee, GA,) and 2 mM L-glut (Life Technologies, cat#35050-061). Propagation of virus stocks was performed in 150 cm^2^ tissue culture flasks. NCTC-1469 cells were infected with stock MHV-A59 at MOI of 0.01 in 10 mL DMEM plus 2% heat inactivated horse serum for 1 hr at 37°C, at which point flasks were supplemented with an additional 8 mL of growth media and cultured at 37°C and 5% CO_2_ until cytopathic effects and syncytia were observed (24 hrs post infection, hpi). Infected cells were then frozen in their flasks at -80°C for 24 hrs, then thawed in a water bath to allow viral release, at which point supernatant was harvested and centrifuged at 1300 xg for 10 mins at 4°C. Viral supernatant was then collected and ultracentrifuged for 3 hrs at 110,500 xg (30,000 rpm in SW32Ti rotor). Virus pellet was resuspended in 1 mL TNE (10 mM TrisCl, 150 mM NaCl, 1 mM EDTA), aliquoted and stored at –80°C until use. Viral titer was determined by plaque assay on L929 (CCL-1) cells obtained from ATCC (Manassas, VA, USA).

### Mice and Infections

8 week old (8 wk) C57BL/6 mice were purchased from Charles River Laboratories (Wilmington, MA) and 18 month old (18 mo) aged mice were obtained from the National Institute on Aging Aged Rodent Colony, also housed at Charles River Laboratories. This study used exclusively male mice. For MHV-A59 infections, mice were deeply anesthetized with isoflurane, then infected i.n. with 10^3^ or 10^4^ pfu MHV-A59 in 50 uL Hanks Balanced Salt Solution (HBSS, Life Technologies, cat#14185052) plus 1% fetal bovine serum (FBS, Life Technologies, cat#13140071). Mock infected controls received only 50 ul inoculum of HBSS plus 1% FBS. All animal experiments were approved by the Institutional Animal Care and Use Committee of the University of North Carolina at Charlotte.

### Viral Burden Measurements

For detection of viral burden, mice were euthanized on designated DPI and blood collected via cardiac puncture into serum separator tubes. After collection, tubes were centrifuged at 1300 rpm for 10 min at 4°C, then serum was transferred into a new microcentrifuge tube. Animals were then perfused with 20 mL cold 1X Phosphate Buffered Saline (PBS, Life Technologies, cat#14200166) and lung, CLN, MdLN, and spleen were collected. Brain was then harvested and microdissected. All organs were snap frozen and weighed. Upon thawing, tissues were homogenized in 500 µL sterile PBS using a bead beater at 4 m/s for 1 min. Viral titers for tissues were assessed by plaque assay on L929 cells. Viral RNA in serum was determined by RT-PCR on RNA collected using E.Z.N.A. Viral RNA Kit (OMEGA Bio-Tek, cat #R6874-01, Norcross GA, USA). Viral genome was quantified using a standard curve of known viral titer following qRT-PCR with iTaq Universal SYBR Green One-Step Kit (BioRad, cat#172-5151, Hercules, CA) and the following primers: Forward: 5’-CAGATCCTTGATGATGGCGTAGT-3’; Reverse: 5’-AGAGTGTCCTATCCCGACTTTCTC-3’ as previously reported(Yang et al., 2014).

### Behavior Testing

Anxiety and baseline locomotor function was assessed using the Open Field assessment. A standard Open Field arena was used consisting of a square box (54 cm x 54 cm) with a grid (6 squares along each side) along the base. Animals were placed into the arena and allowed to explore for 5 minutes, at which point the animal was returned to its home cage and the arena sanitized with 70% ethanol between trials. Trials were recorded using Stoelting ANY-maze USB 2 camera (Stoelting Co; Wood Dale, IL) and the number of lines crossed, total time spent in the center of the arena, and average speed was measured using ANY-maze software (Stoelting Co; Wood Dale, IL). Spatial memory was assessed using the Barnes Maze. An elevated Barnes Maze table (91.4 cm diameter, containing 19 empty holes and 1 target hole, spaced evenly 5 cm apart and 6.25 cm from the edge of the table) was used for testing. Mice were tested over 5 consecutive days, with 2 trials each day, spaced exactly 30 min apart. Visual cues around the room were retained in the same location for the duration of the testing period. For each trial, the animal was placed into the center of the maze in a covered start box for 10 s, with removal of the box signaling start of the trial. Each animal was given 3 min to explore the maze and find the target hole. Animals that did not locate the target hole within 3 min were gently guided into it. After each trial, the animal remained in the target hole for exactly 1 min, then was returned to their home cage. The maze was decontaminated with 70% ethanol between trials. Trials were recorded using Stoelting ANY-maze USB 2 camera (Stoelting Co; Wood Dale, IL) and latency to find the target hole was determined using ANY-maze software (Stoelting Co; Wood Dale, IL).

### Tissue Isolation for Flow Cytometry

Single cell suspension from tissues was performed as previously described(Shane et al., 2018). Briefly, after perfusion with 20 mL PBS, lungs were excised, minced and incubated for 30 min at 37°C with 1.25 mM EDTA, followed by 1 hr incubation with 150 U/mL collagenase (Life Technologies, cat#17-104-019)) diluted in RPMI (Life Technologies, cat#21-870-100) supplemented with 1.25 mM CaCl_2_ (Fisher, cat#AC349610250, Waltham, MA), 1.25 mM MgCl_2_ (Fisher, cat#AC223211000), 5% FBS, 500 mM HEPES (Life Technologies, cat# 15630080), 200 mM L-glut (Life Technologies, cat#21-051-024), 2000 U/mL Antibiotic-Antimycotic (Fisher, cat#15240062), and 5 ug/mL Gentamycin (Life Technologies, cat#15750060). Cells were then passed through a 40 µM cell strainer and resuspended in 44% Percoll (Fisher, cat#45-001-747) diluted in PBS underlaid with 67% Percoll diluted in RPMI, then centrifuged at 2800 rpm for 20 min at 4°C and the cellular interface collected. For isolation of lymphocytes from the brain tissue, perfused brains were excised, minced, and incubated for 1 hr with 5 mg/mL collagenase type I (Sigma, cat#C0130-500MG, St Louis, MO) diluted in HBSS supplemented with 0.1 ug/mL TLCK trypsin inhibitor (Sigma, cat#T754), 10 ug/mL DNase1 (Sigma, cat#D4263-0.5MG), and 10 mM HEPES (Life Technologies, cat# 15630080). Cells were then passed through a 40 µM cell strainer and resuspended in 37% Percoll diluted in RPMI, centrifuged at 2800 rpm for 20 min at 4°C and cell pellet collected. Lymph nodes and spleens were mechanically digested by passage through a 40 µM cell strainer and collected in RPMI. Erythrocytes were removed from spleens using ACK lysis buffer (Life Technologies, cat#A1049201). Live cell numbers of all samples were determined by staining with Trypan blue (Life Technologies, cat#15250061) and counted using Bio-Rad TC-20 Automated Cell Counter (Hercules, CA).

### Flow Cytometry

Prior to immunostaining, all cells were blocked with 1:50 dilution of TruStain FcX anti-mouse CD16/CD32 (Clone 93, Biolegend) and Fixable Viability Dye (eBioscience, cat#65-0866-18, San Diego, CA) for 5 mins. The MHV spike protein (S) MHC class I tetramer [H-2K^b^/RCQIFANI] was generated at the National Institute of Allergy and Infectious Diseases Tetramer Facility (Emory University, Atlanta, GA). Tetramer staining was carried out at room temperature for 20 minutes in conjugation with other surface staining antibodies: CD8a (53-6.7, APC-Cy7), CD4 (RM4-5, APC), CD44 (IM7, PE-Cy7), CD45 (30-F11, APC), CD11b (M1/70, FITC), P2RY12 (S16007D, PE), CD68 (FA-11, BV-421), MHC II (M5/114.15.2, PerCP-Cy5.5), obtained from Cytek Biosciences (Fremont, CA) or eBioscience (San Diego, CA), then washed with 1X PBS and fixed with 2% paraformaldehyde (PFA). Data was acquired using a BD Fortessa with FACSDiva Software (BD Biosciences, San Jose, CA) and analyzed using FlowJo software version 10.9.0 (BD Biosciences, San Jose, CA). All samples were gated on live, single cells, followed by lymphocytes as determined by forward (FSC-A) and side scatter (SSC-A) unless otherwise indicated.

For assessment of intracellular cytokine production, cells were harvested as described above and treated with eBioscience Cell Stimulation Cocktail (cat#00-4970-03) and eBioscience Protein Transport Inhibitor (cat#00-4980-03) for 5 hrs at 37°C. Following stimulation, cells were stained for cell surface markers, then permeabilized using eBioscience PermWash and intracellularly stained as per the manufacturer instructions (cat#00-5523-00) for IFN-γ (XMG1.2, APC), TNF (MP6-XT22, PerCP-710), perforin (eBio0MAK-D, FITC), and granzyme B (NGZB, PE-Cy7). Cells were then fixed in 2% paraformaldehyde for flow cytometry analysis.

### Immunohistochemistry

At designated timepoints post infection, animals were euthanized and perfused with 20 mL cold PBS, then brains were excised and fixed overnight in 4% PFA. Following fixation brains were cryoprotected in two exchanges of 30% sucrose diluted in PBS, then embedded and frozen in OCT (Fisher, cat#23-730-571). Sagittal sections were cut to 10 µm using a Microm HM550 cryostat. Prior to immunostaining, sections were blocked for 1 hr at room temperature with 5% goat serum/0.1% Tween-20/PBS, then stained at 4°C overnight with various combinations of the following antibodies: rabbit anti-NeuN (1:1000, Cell Signaling Technologies, 12943S), rabbit anti-Iba1 (1:1000, Wako Chemicals, 019-19741), rabbit anti-GFAP (1:1000, Cell Signaling Technologies, 12389S), guinea pig anti-synaptophysin (1:500, Cedar Lane Labs, 101004(SY)), guinea pig anti-DCX (1:500, MilliporeSigma, AB2253 ), mouse anti-MHV Nucleocapsid (1:1000, BEI, NR-45106). Sections were then washed with 0.1% Tween-20/PBS and incubated for 1 hr at room temperature with the following fluorescently labeled secondary antibodies: Alexa Fluor 647 anti-Rabbit (1:400, Life Technologies, cat#A-21245), Alexa Fluor 488 anti-Guinea-Pig (1:400, Life Technologies, cat#A-11073), Alexa Fluor 488 anti-Rat (1:400, Life Technologies, cat#A-11006). Sections were then counterstained with 1 µg/mL DAPI and coverslipped with Prolong Gold Antifade Mountant (Fisher, cat#P36930). Samples were treated with TUNEL staining reagent as prepared by manufacturer’s instructions (Roche in situ cell death kit, TMR Red, cat#12-156-792-910) prior to blocking and primary staining. Images were captured using a Keyence BZX-800 at 20x and 40x objective magnification. Images were processed and quantified using Fiji-ImageJ(Schindelin et al., 2009).

### In vitro CD8^+^ T cell Neuron Co-cultures

Primary cortical neurons were isolated from embryos collected from timed-pregnant C57BL/6 mice on embryonic day 18 as previously described(Funk and Lotz, 2020). Briefly, cerebral cortices were dissected from embryonic brains, treated with papain (20 U/mL) and DNase 1 (2.5 U/mL) for 30 minutes, and gently dissociated by trituration in Hibernate E medium (Life Technologies, cat#A1247601). Neurons were diluted in 5 ml Neuron Growth Medium (Neurobasal media (Life Technologies, cat#A3582901) supplemented with 2% B27 (Life Technologies, cat#17504001), 2 mM L-glut (Life Technologies, cat#35050-061) and 100 U/mL Antibiotic-Antimycotic (Fisher, cat#15240062)) at a concentration of 1x10^5^ neurons/ml so that 1x10^5^ neurons were seeded onto poly-D-lysine (10 µg/mL) coated acid-etched coverslips (VWR,, cat#MSPP-P06G1520F) and cultured at 37°C plus 5% CO_2_. Half-media changes were performed every 72 hrs until neurons were used for experiments at 7-10 days *in vitro*, as indicated in text.

CD8^+^ T cells were isolated from the spleens of uninfected or MHV-A59 infected animals using the untouched CD8^+^ T cell isolation MACS kit from Miltenyi Biotec (cat#130-104-075, Gaithersburg, MD) according to manufacturer’s instructions. CD8^+^ T cells were seeded onto neurons at a concentration of 1x10^5^ cells/well (1:1 ratio) for 24 hrs, at which point culture supernatant was collected and cryopreserved, and coverslips were co-stained for TUNEL and NeuN, as described above. Neuronal cytotoxicity was also examined at 24 hrs post culture using the Promega CellTiter Blue Cell Viability Kit (Fisher, cat#PRG8080) according to the manufacturer’s instructions.

### ELISA

IFN-γ protein levels were measured from cryopreserved neuron-CD8^+^ T cell co-culture supernatant using the Mouse IFN-γ DuoSet ELISA kit from R&D Systems (R&D Systems, cat#DY48505, Minneapolis, MN) according to the manufacturer’s instructions.

### Statistics

Statistical analysis was performed using Graph Pad Prism 10.0.2.

## Data Availability Statement

All data are available in the article and its supplemental material.

## Acknowledgements

This work was supported by National Institutes of Health grant R00AG053412 and R00AG053412-04S1. We thank the NIH Tetramer Core Facility (contract number 75N93020D00005) for providing MHV-S tetramers and the NIA Aged Rodent Colony for providing aged mice used in this this study. The authors declare no competing financial interests.

## Abbreviations

BBB: Blood-brain barrier
CLN: Cervical lymph nodes
CNS: Central nervous system
DCX: Doublecortin
DPI: Days post-infection
FSC: Forward scatter
I.N.: Intranasal
MBP: Myelin basic protein
MDLN: Mediastinal lymph nodes
MHV-A59: mouse hepatitis virus strain A59
MOI: Multiplicity of infection
PFU: Plaque-forming units
SSC: Side scatter
WNV: West Nile virus

