## Supplemental Figures and Legends for "Antigen non-specific CD8^+^ T cells accelerate cognitive decline in aged mice following respiratory coronavirus infection"

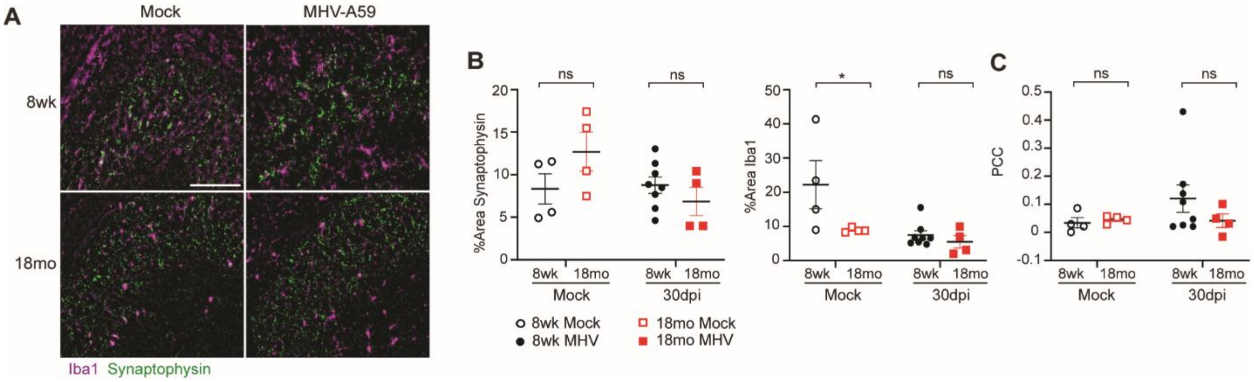


**Supplemental Figure 1.** Assessment of microglia mediated synaptic elimination following MHV-A59 infection.

(**A**) Representative IHC images of Iba1 and Synaptophysin in the CA3 region of the hippocampus of 8 wk or 18 mo old animals at 30 DPI. Images taken at 40X magnification. Scale bar = 100 µm. (**B**) Quantification of percent area of Synaptophysin and Iba1 staining. (**C**) Quantification of PCC for synaptophysin and Iba1 colocalization.

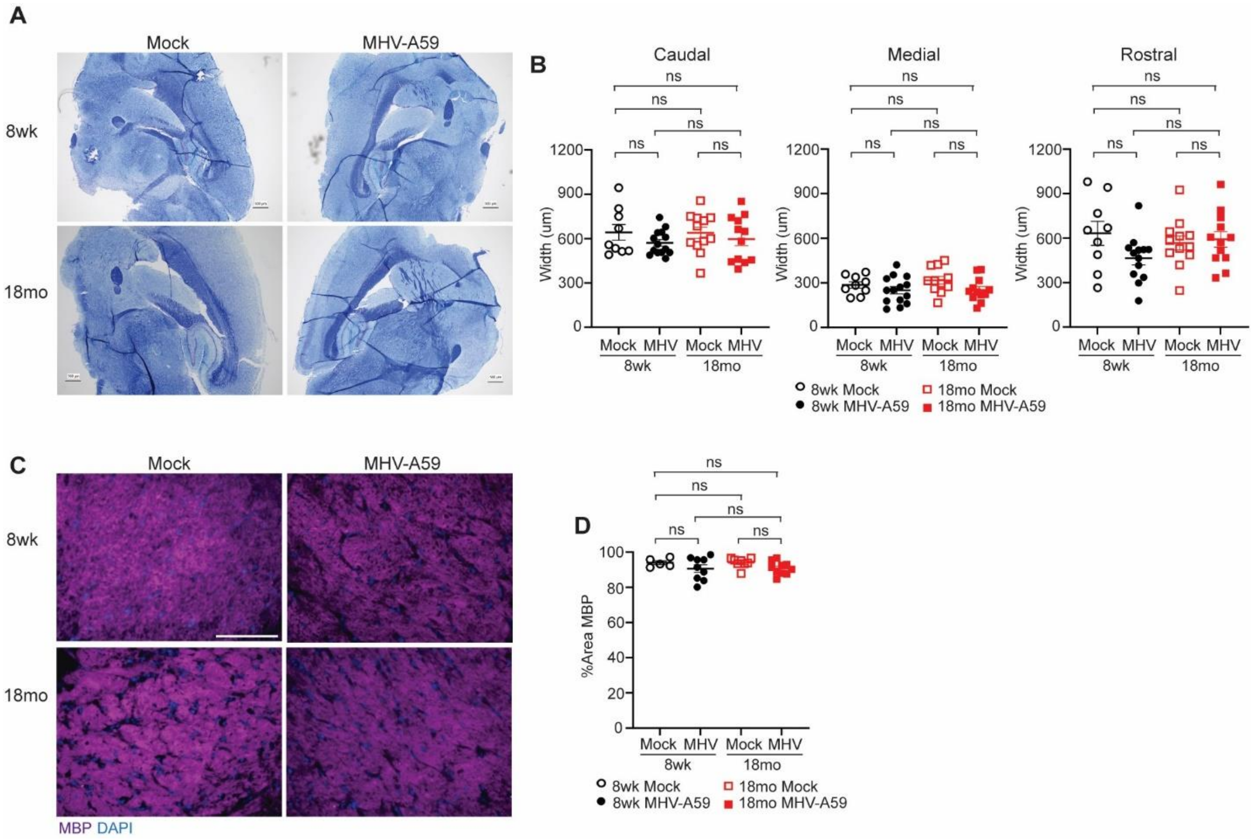


**Supplemental Figure 2.** Assessment of demyelination in adult and aged animals 30 DPI MHV-A59 infection.

(**A)** Representative brightfield images of Luxol Fast Blue staining in the corpus callosum of 8 wk or 18 mo old MHV-A59 or mock infected animals 30 DPI. Images taken at 2X magnification. Scale bar = 500 µm. (**B**) Quantification of caudal, medial, and rostral corpus callosum width. (**C**) Representative IHC images of MBP and DAPI staining within the corpus callosum of 8 wk or 18 mo old MHV-A59 or mock infected animals 30 DPI. Images taken at 40X magnification. Scale bar = 100 µm. (**D**) Quantification of percent area of MBP in the corpus callosum. Images taken at 40X magnification. Scale bar = 100 µm. Statistics according to unpaired one-way ANOVA conducted in Graph Pad prism 9.3.1 software. *, $p<0.05$

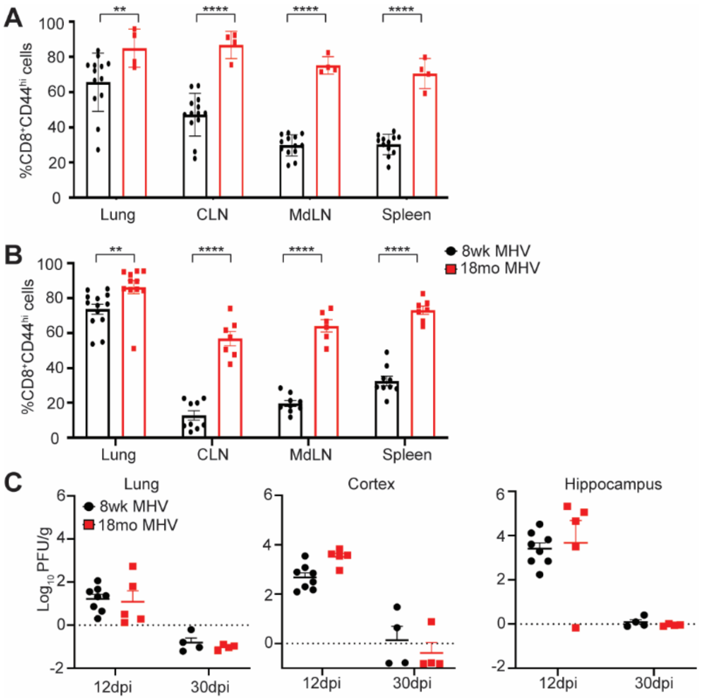


**Supplemental Figure 3.** Enhanced CD8$^+$ T cells responses in peripheral compartments of aged mice following MHV-A59 infection

8 wk adult or 18 mo aged C57BL/6 mice were infected with (**A**) $10^4$ pfu MHV-A59 or (**B**) $10^3$ pfu MHV-A59, and frequency of CD8$^+$CD44$^{hi}$ cells present at 30 DPI were quantified by flow cytometry. Data were pooled from 2 independent experiments with each data point representing a single animal. (**C**) Viral genome was determined by qRT-PCR. Statistics according to two-way ANOVA conducted in Graph Pad prism 9.3.1 software. *, $p<0.05$; **, $p<0.01$; ***, $p<0.001$; ****, $p<0.0001$.

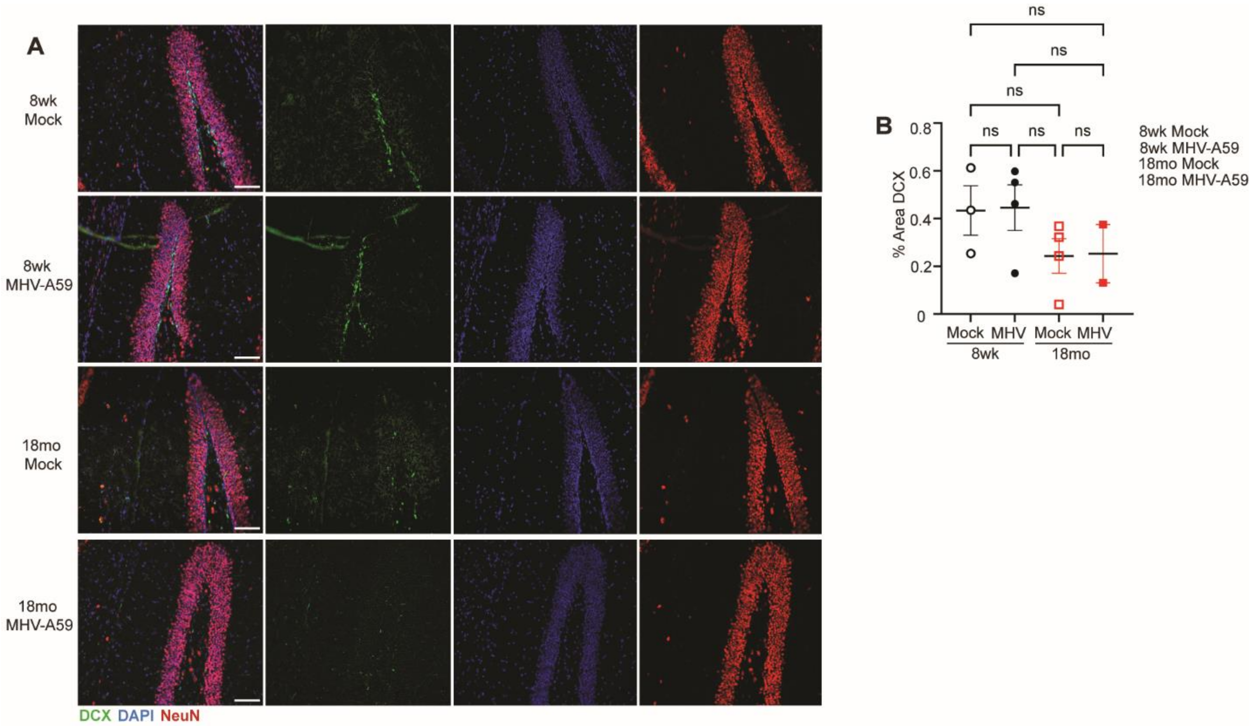


**Supplemental Figure 4.** Impaired neuronal regeneration in aged animals.

(**A**) Representative IHC of DAPI, NeuN, and DCX in the DG of 8 wk or 18 mo old animals at 30 DPI compared to mock infected controls. Images taken at 20X magnification. Scale bar = 100 µm. (**B**) Quantification of total percent area of DCX in the DG. Data representative of 1 experiment with each data point representing a single animal. Statistics according to unpaired one-way ANOVA conducted in Graph Pad prism 9.3.1 software. *, $p<0.05$; **, $p<0.01$; ***, $p<0.001$; ****, $p<0.0001$.

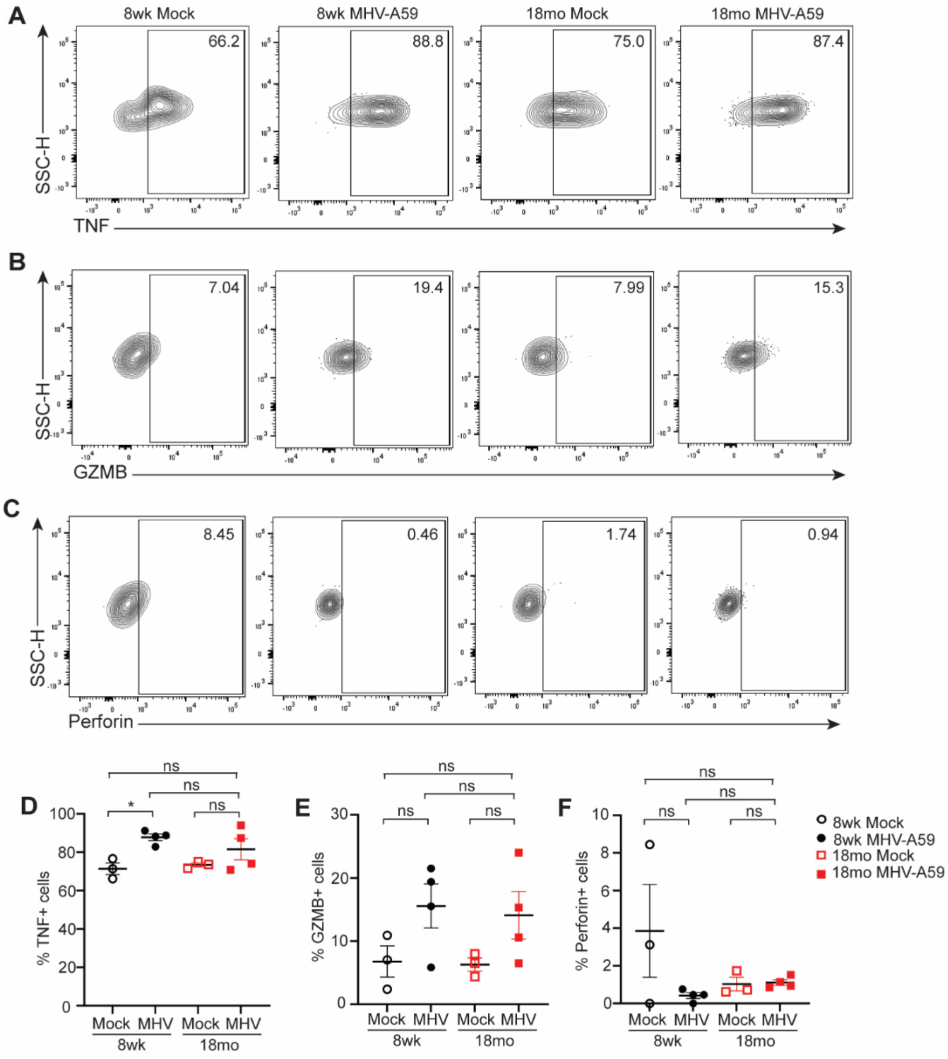


**Supplemental Figure 5.** Production of TNF, GZMB, and Perforin by 8 wk or 18 mo old animals at 30 DPI

(**A-C**) Representative flow cytometry plots of (**A**) $TNF^{+}$, (**B**) $GZMB^{+}$, or (**C**) $Perforin^{+}$ $CD8^{+}$ T cells isolated from the brain of 8wk or 18month old MHV-A59 or mock infected animals 30 DPI following 4 hr PMA/ionomycin stimulation. (**D-F**) Quantification of percent frequency of $CD8^{+}$ $CD44^{+}$ lymphocytes expressing (**D**) TNF, (**E**) GZMB, or (**F**) Perforin. Cells previously gated on live, single cell, $CD8^{+}$ $CD44^{+}$ lymphocytes. Statistics according to unpaired one-way ANOVA conducted in Graph Pad prism 9.3.1 software. *, $p<0.05$
